# Scaling read aligners to hundreds of threads on general-purpose processors

**DOI:** 10.1101/205328

**Authors:** Ben Langmead, Christopher Wilks, Valentin Antonescu, Rone Charles

## Abstract

General-purpose processors can now contain many dozens of processor cores and support hundreds of simultaneous threads of execution. To make best use of these threads, genomics software must contend with new and subtle computer architecture issues. We discuss some of these and propose methods for improving thread scaling in tools that analyze each read independently, such as read aligners. We implement these methods in new versions of Bowtie, Bowtie 2 and HISAT. We greatly improve thread scaling in many scenarios, including on the recent Intel Xeon Phi architecture. We also highlight how bottlenecks are exacerbated by variable-record-length file formats like FASTQ and suggest changes that enable superior scaling.

## 1 Introduction

General-purpose processors are now capable of running hundreds of threads of execution simultaneously in parallel. Intel’s Xeon Phi “Knight’s Landing” architecture supports 256-288 simultaneous threads across 64-72 physical processor cores [1, 2]. With severe physical limits on clock speed [3], future architectures will likely support more simultaneous threads rather than faster individual cores [4]. Indeed, clock speed on the many-core Xeon Phi processor (1.3-1.5 Ghz) is about half that of more typical server processors. While specialized (e.g. graphics) processors have been highly multithreaded for some time, this only recently became true for the general-purpose processors that can boot standard operating systems and that typically power servers and desktops.

With these advances come new computer-architecture considerations for programmers. Simply adding multi-threading to a software tool does not guarantee it will use threads well. In fact, it is not uncommon for a tool’s overall throughput to *decrease* when thread count grows large enough [5]. So whereas past genomics software efforts have focused on speed on a fixed (and usually low) number of threads, future efforts should consider scaling to much higher thread counts.

Here we tackle the problem of scaling read aligners to hundreds of threads on general-purpose processors. We concentrate on the Bowtie [6], Bowtie 2 [7] and HISAT [8] read alignment tools because they are widely used and representative of a wider group of *embarrassingly parallel* tools, where computation is readily separable into independent tasks, one per sequencing read. Many other sequencing analysis tools are also embarrassingly parallel, including tools for error correction [9, 10], quality assessment and trimming [11], and taxonomic assignment [12, 13].

We propose strategies that scale to hundreds of threads better than alternative approaches like multiprocessing or the pipelined approach taken by BWA-MEM [14]. We explore how the FASTQ file format [15], its unpredictable record boundaries in particular, can impede thread scaling. We suggest a way to change FASTQ files and similar formats that enable further improvements in thread scaling while maintaining essentially the same compressed file size.

### Synchronization and locking

For embarrassingly parallel genomics tools, threads typically proceed by repeatedly (a) obtaining the next read from the input file, (b) aligning the read, and (c) writing its alignment to the output file. Interactions with input and output files must be synchronized; portions of code related to reading and writing files must be protected to allow only one thread at a time to work on a given file. Figure 1 illustrates threads operating in parallel while reading input in a synchronized fashion. Failure to synchronize can lead to software crashes and corrupt data. Synchronization is achieved with locks. There are various lock types, which incur different types and amounts of overhead. We confirm here that for many-core architectures with non-uniform memory access (NUMA), choice of lock type has a major impact on thread scaling [16]. We explore several lock types, demonstrate their relative merits, and suggest types to be avoided (spin locks) and others that seem to scale well to hundreds of threads (queueing and standard locks).

**Figure 1:**
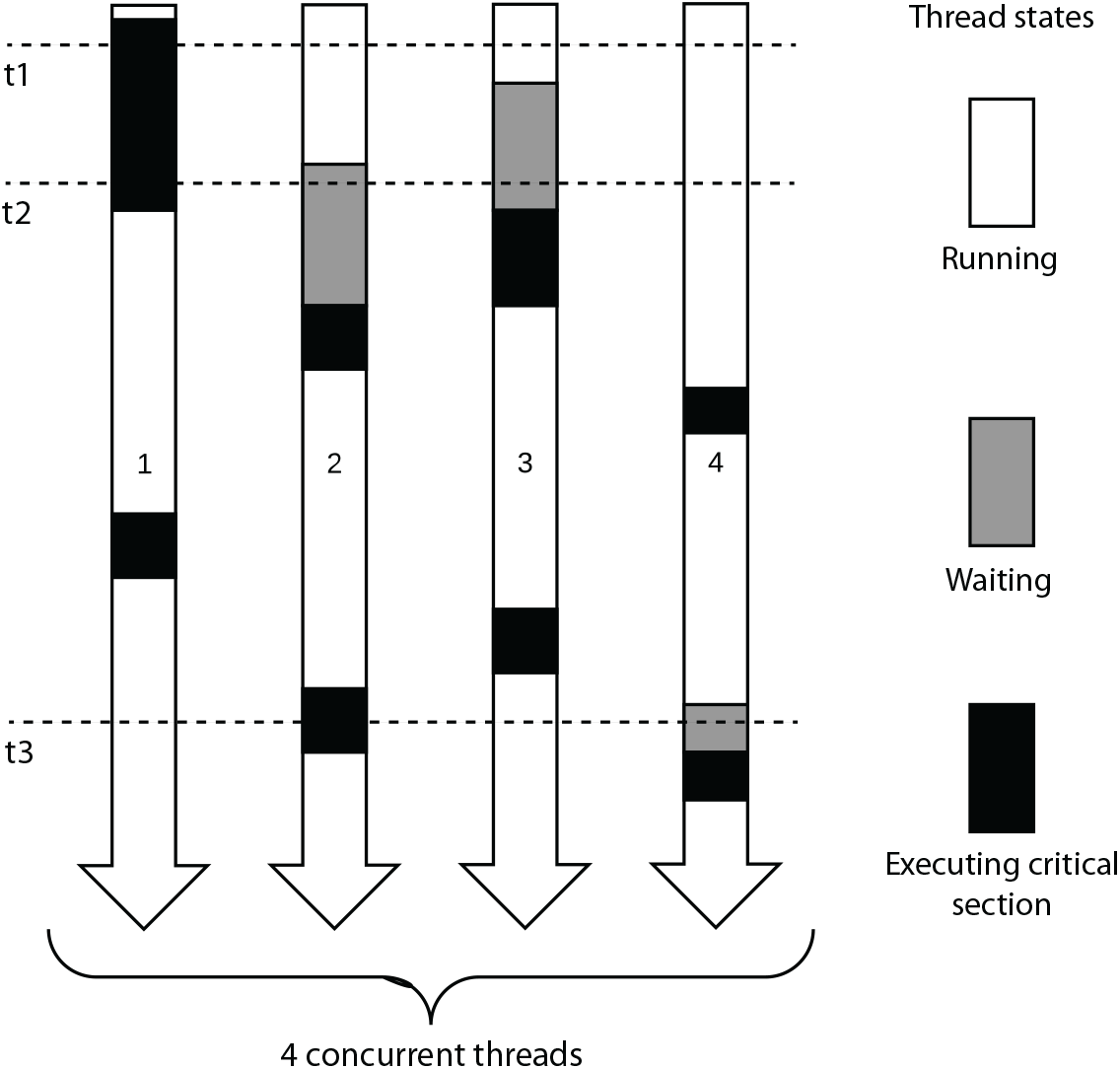
Four threads running simultaneously in an embarrassingly parallel setting. Time progresses from top to bottom. Gray boxes show time spent waiting to enter the critical section. Black boxes show time spent in the critical section, which can be occupied by at most one thread at a time. At time t1 (dashed line), thread 1 is executing the critical section and all the other threads are running. At time t2, thread 1 is still in the critical section and threads 2 and 3 are waiting to enter. At time t3, thread 2 occupies the critical section and thread 4 is waiting.

### Multithreading versus multiprocessing

While we focus on making the best use of threads in a single process, an alternate is to run multiple simultaneous processes, possibly with many threads each. For example, a user with several FASTQ files might align all using Bowtie 2 with the `-p 100` argument, using 1 process with 100 threads. Alternately, the user could divide the input into 10 batches and run 10 simultaneous Bowtie 2 processes each with `-p 10`. Either way, up to 100 threads run in parallel.

These multithreading (MT) and multiprocessing (MP) approaches have trade-offs. MP can suffer from *load imbalance*: some batches take longer to align than others. This negatively impacts scaling since the job’s duration is determined by the longest-running batch.

Imbalance can be mitigated by *dynamic* load balancing. Such a scheme might divide the input into many batches, more than there are processes. A load balancer launches the processes and continually feed each process new input batches upon completion of the previous batch, until all batches are processed. For a large enough number of batches, per-batch running times tend to average out, making per-process running times more uniform. This incurs overhead, since the dynamic load balancer must split inputs, launch and feed processes, and combine outputs.

Another drawback of MP is that many data structures will be copied across processes, yielding a higher total memory footprint compared to MT. When the data structure is identical from process to process (e.g. the genome index), this wastes valuable memory and cache. Aligners with large indexes, e.g. GEM [17] and SNAP [18], can easily exhaust available memory [5]. Tools can work around this by using memory mapping to maintain a single copy of these data structures shared by all processes. This is implemented in Bowtie, Bowtie 2 and HISAT using the `--mm` option.

The MP strategy also has a major advantage: by allowing each processes to focus on its own private input and output files, the overall level of thread contention is reduced. That is, a single process has fewer threads to synchronize. There are also NUMA-related reasons why running multiple processes can aid thread scaling, e.g. by allowing each process to have a copy of the genome index that is local to its home NUMA node [5].

The MT approach has many advantages. It achieves dynamic load balancing without extra software beyond the aligner itself. It achieves a low memory footprint (no duplicated data structures) without the need for memory mapping. It is applicable regardless of the number of input files, naturally handling the common case of a single, large input file. For these reasons, we focus on improving MT thread scaling in this study, using MP as a baseline.

### Input and output

Improving speed and thread scaling can eventually reach a point where the bottleneck shifts from the speed of computation to the speed of input and/or output. Since we would like to observe whether this occurs, all our experiments use real input and output. As discussed later, while input and output speed are not bottlenecks for most of our experiments, there are scenarios where output becomes the bottleneck on very large numbers of threads.

### Related work

Two prior studies [19, 20], examined Bowtie 2 thread scaling with synchronization and Non-Uniform Memory Access (NUMA) as primary concerns. By adapting Bowtie 2 to the FastFlow [21] parallel framework and by making effective use of (a) thread pinning and (b) interleaving of memory pages across NUMA sockets, the modifications improved Bowtie 2’s thread scaling. Our suggestions for improving thread scaling are complementary to these proposals.

Herzeel et al [22] re-parallelized sections of the BWA code using the Cilk [23] programming language. They noted a 2-fold improvement in multi-threaded speedup, highlighting the importance of NUMA and load balance issues. Lenis and Senar examined performance of four read aligners, including Bowtie 2 and BWA-MEM on NUMA architectures [5] without modifications, and noted that a multiprocessing approach that replicated the index data structure across NUMA nodes performed the best. Our goal is to achieve similar improvements with a purely multithreaded approach on modern hardware.

## 2 Methods

### 2.1 Lock types

We begin by examining how lock types affect thread scaling. Different lock types are appropriate for different situations. A *spinlock* uses a loop to repeatedly check if a lock is held. As soon as a check indicates the lock is free, ownership is transferred to the inquiring thread. The check and the transfer can happen simultaneously using an atomic operation [24]. In most implementations, a thread that fails to obtain a spinlock in a prescribed time interval will go to sleep, allowing the operating system (OS) to revive it when the lock is free. This avoids *starvation*, whereby the lock-holding thread is slow to finish its work (and release the lock) because waiting threads are using its resources. This, spinlocks are *optimistic*: they work best when the lock can be obtained quickly.

Another common lock type is a *standard lock*; if a thread attempts and fails to obtain a standard lock, it goes to sleep immediately, allowing the OS to revive it when the lock is free. While pausing and reviving a thread incurs overhead, a standard lock cannot starve other threads. Thus, a standard lock is *pessimistic*, working best when the lock is unlikely to be available soon.

We might suppose that when active thread count is less than or equal to the number of physical cores — a typical situation when a user has dedicated access to a computer and desires speed — starvation is not an issue and spinlocks are ideal. However, this supposition fails on modern many-core systems for reasons relevant to our choice of lock type. One concern is that modern architectures have many cores and caches connected in a NUMA architecture. That is, there is a single addressable memory space for all threads, but it is physically divided into partitions that might be attached to separate cores in a multi-socket system, as for the 2-socket Broadwell system used in our evaluations, or both the partitions and the cores might be connected via an interconnection network, as on the Xeon Phi. Thread scaling is impacted in at least two ways: (a) threads using different cores but accessing the same memory location will incur different access latencies depending on the distance to the memory, and (b) when several threads read and write the same location simultaneously, the system’s *cache coherence protocol* must step in to ensure all threads have a coherent view of memory. In short, thread scaling suffers when added threads must access distant memories or when they compete for the same memory locations as existing threads. This affects locking in key ways that we revisit when discussing the queueing lock.

Another issue arises when threads can co-exist on the same physical processor. On Xeon Phi, up to four threads can run simultaneously on one processor, competing for its resources like its arithmetic units and cache. A thread operating by itself on a processor moves at one speed, but slows when joined by a second thread, slows still further when joined by third, etc. Thus, increasing thread count incurs a mild but increasing starvation penalty even when free thread “slots” remain. This puts optimistic locks at a disadvantage, since their spinning behavior can needlessly starve productive threads on the same processor.

In past versions, Bowtie, Bowtie 2 and HISAT used a spinlock from the `TinyThread++` library (http://tinythreadpp.bitsnbites.eu). Since this scaled poorly (see Results), we extended the three tools to use the open source Intel Thread Building Blocks (TBB) library [25]. TBB provides various lock types, including a queuing lock [26] particularly appropriate for NUMA systems like the Knight’s Landing and Broadwell systems used here. TBB also provides scalable replacements for standard heap memory allocation functions (e.g. `malloc/free, new/delete`). This aids thread scaling, since memory allocations require synchronization.

The TBB queuing lock implements an MCS lock [26], which uses an in-memory queue to organize waiting threads. Like a spinlock, a waiting thread repeatedly probes a variable in memory to learn when it has obtained the lock. Unlike a spinlock, each waiting thread probes a separate queue entry, each entry occupying a separate cache line. This greatly reduces overhead. To elaborate, consider that an atomic operation (e.g. atomic compare-and-swap) might modify a variable in memory, depending on the condition. Consequently, it is treated as a memory write by the cache coherence infrastructure. A write modifies a cache line, causing cache coherence messages to travel between caches for threads that recently accessed the line. When this happens in a loop, new messages are generated each iteration. When many threads spin simultaneously, messages multiply, eventually reaching a point where the messages flood the system bus and starve other threads, including lock holder. This is called *cache-line* or *hotspot* contention [26] and it is a major concern on many-core and NUMA systems [16]. The queuing lock reduces contention in two ways. First, since each thread spins on a variable in a thread-specific cache line, the loop condition can be a simple memory read rather than an atomic operation. This reduces cache coherence messaging. Second, while a memory write is still needed to hand the lock from one thread to another, only two threads are involved in the hand-off, reducing the coherence messages exchanged.

We adapted the three tools to use four lock types: the (original) `TinyThread++` lock, standard TBB lock, TBB spinlock, and TBB queuing lock. On the Linux systems we used for evaluation, the standard TBB lock works by calling `pthread_mutex_unlock`, which in turn uses the Linux `futex` (fast mutex) strategy. This strategy first attempts to obtain the lock using a fast atomic operation then, if unsuccessful (i.e. if the lock is held by another thread), places it on a queue of paused threads until the lock is released.

The lock type is selected at compile time via preprocessing macros. These extensions are available as of the Bowtie v1.1.2, Bowtie 2 v2.2.9 and HISAT v0.1.6-beta software versions. Supplementary Note 1 gives build instructions for the exact software versions tested here.

### 2.2 Parsing strategies

We also examined how threads coordinate when reading FASTQ [15] input or writing SAM output [27]. These interactions are synchronized, i.e. protected by locks. The name *critical section* is given to a portion of the software that only one thread may execute at a time. The critical section for handling input is called the *input critical section* and is protected by the *input lock*; likewise for the *output critical section* and the *output lock*.

We hypothesized that to improve thread scaling we should restructure the input and output critical sections. Our first goal was to reduce the time spent in the critical section by deferring as much computation until after the critical section as possible. Our second goal was to reduce the total number of times the critical section was entered. This reduces overhead incurred by locking and unlocking upon entering and exiting.

The original strategy (O-parsing) both reads and parses a sequencing read in the critical section (CS). We developed three variants on this approach (Table 1). In deferred (D) parsing, the CS reads a single input record into a buffer. After the CS, the buffer is parsed into the sequencing read data object. Batch deferred (B) parsing is like D-parsing but handles batches of *N* reads at a time. The B-parsing critical section loops *N* times, reading each record into a separate buffer. After the CS, another loop parses each buffer into a sequencing read object. This reduces by a factor of *N* the total number of times the CS is entered. A similar change is made to the output CS: alignment records are written to the output stream in batches of *N* reads.

**Table 1:** Pseudocode for four synchronized parsing strategies. Red code is inside the critical section (CS). Original (O) parsing both reads and parses in the CS. Deferred parsing (D) uses the CS to read the next record into a buffer, counting four newlines to find the record boundary, but defers parsing until after the CS. Batch deferred parsing (B) is like (D) but reads N reads at a time. Block deferred parsing (L) reads a fixed-sized chunk of data (*B* bytes), assuming that no record spans a *B*-byte boundary. While the assumption for (L) is violated in practice for formats like FASTQ, it suggests a strategy for making formats more amenable to multithreaded parsing.

| (Original) O-parsing | (Deferred) D-parsing |
| --- | --- |
| `obtain lock read = read_and_parse() release lock` | `buf = new empty buffer obtain lock nl = 0 while nl < 4: c = read_char() if c is newline: nl = nl + 1 append c to buf release lock read = parse_fastq(buf)` |
| (Batch deferred) B-parsing | (Block deferred) L-parsing |
| `bufs = array of N empty buffers reads = array of N empty reads obtain lock for i = 1 to N: nl = 0 while nl < 4: c = read_char() if c is newline: nl = nl + 1 append c to bufs[i] release lock for i = 1 to N: reads[i] = parse_fastq(bufs[i])` | `reads = array of N empty reads obtain lock buf = read B bytes release lock for i = 1 to N: reads[i] = parse_fastq(buf) advance buf to next read` |

Blocked deferred (L) parsing reads a chunk of exactly *B* input bytes into a buffer, assuming that (a) no read spans a *B*-byte boundary in the input file, such that no *B*-byte chunk contains a partial input record, and (b) the number of reads per *B*-byte chunk is *N* for all chunks (except perhaps the last), known ahead of time. These assumptions do not hold for real FASTQ files, but we can easily modify a FASTQ file to comply by appending extra space characters to every *N*th read until the following read begins at an *B*-byte boundary (Figure 2). The spaces are ignored by the aligner. This has the effect both of enforcing the L-parsing assumptions and of making it easier to parse paired-end files in a synchronized manner, since a *B*-sized block taken from the same offset in both files is guaranteed to contain *N* matching ends. As with *B*-parsing, the L-parsing output critical section writes alignments in batches of *B* reads at a time.

**Figure 2:**
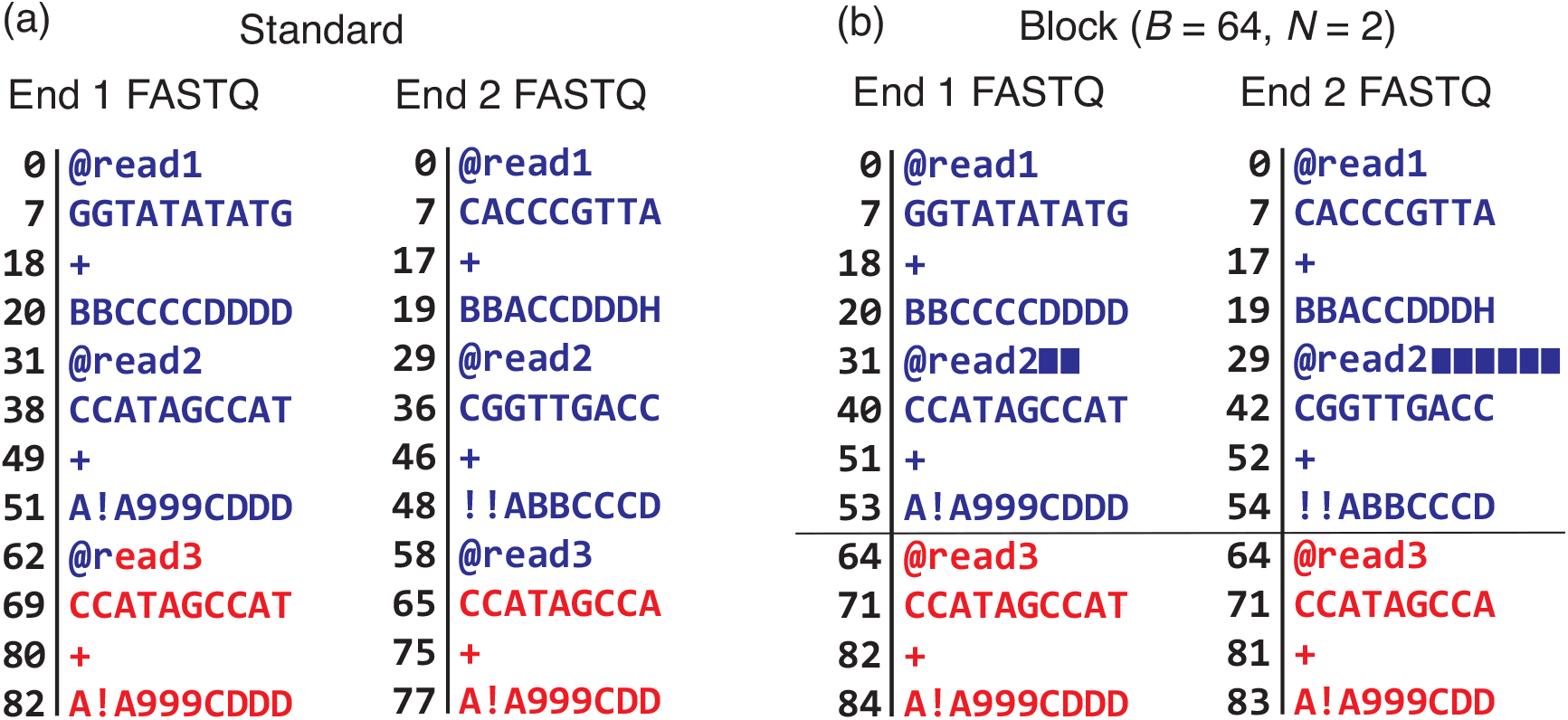
Converting a standard pair of FASTQ files (a) to blocked FASTQ files (b), where the number of bytes (*B*) and number of input read per block (*N*) are 64 and 2 respectively. Numbers left of vertical lines indicate byte offsets for FASTQ lines, assuming newline characters (not shown) are one byte. For (b), padding spaces are represented by solid blue rectangles. The first 64 bytes of each file are colored blue and subsequent bytes are colored red. Note that the two ends differ in length; end 1 is 10 bases long and end 2 is 9 bases long. This necessitates differing amounts of padding in the two FASTQ files. But after padding, we are guaranteed that corresponding 64-byte blocks from the files contain *N* corresponding reads.

### 2.3 Output striping

While most improvements proposed here reduce input synchronization overhead, we also noted instances where output synchronization was the bottleneck. Synchronized output is quite simple; no parsing is involved. But it can still become a bottleneck since writing is generally much slower than reading. On the Stampede 2 cluster, for example, read throughput is about 3 GB/sec when reading from solid-state or Lustre storage, whereas writing is about 450 MB/sec for solid-state and 300 MB/sec for Lustre. To this end, we implemented the ability to write *striped* output, i.e. multiple output SAM files, each containing alignment for a subset of the reads. The partial output files can simply be concatenated prior to further processing. This spreads contention for the output lock over several locks, reducing contention and improving scalability.

### 2.4 Other aligner modifications

We also modified the aligners to minimize the incidence of heap memory allocations wherever possible. This is because heap memory allocations also require synchronization, thus negatively impacting thread scaling. We also modified each of Bowtie, Bowtie 2 and HISAT to cause each thread to report how much wall-clock time it spends aligning reads with microsecond accuracy.

### 2.5 Multiprocessing

While our focus is on single-process multithreaded (MT) approaches, multiprocessing (MP) is another avenue for improving thread scaling. For this reason, our comparisons include an MP-based “baseline” strategy. The MP baseline is measured for every thread count *T* that is multiple of 16 by running T/16 processes, each with 16 threads. For MP experiments involving Bowtie, Bowtie 2 or HISAT, we use memory mapping (`--mm` option) to limit overall memory footprint.

## 3 Results

### Configurations & jobs

We evaluate various read aligners and synchronization schemes by running each “configuration” (combination of aligner and synchronization scheme) using the same input data. For each configuration, we perform a series of alignment *jobs* varying the number of input reads and the number of simultaneous threads of execution in direct proportion, thus keeping the number of reads per thread constant. In this way, we are assessing *weak scaling*: how running time varies with the number of threads for fixed *per-thread* workload.

We align to an index of the GRCh38 human genome reference assembly [28]. We measure wall-clock running time of each job, omitting time required for one-time setup tasks such as index loading, since these influence thread scaling only slightly when aligning large datasets. The number of reads per thread (Supplementary Table 1) was chosen for each configuration and system so that most jobs take 1 minute or longer. Any job taking longer than 20 minutes was aborted and omitted from the results. The Linux top utility was run in the background to periodically measure system load, processor utilization and memory footprint.

We evaluate Bowtie [6], Bowtie 2 [7] and HISAT [8] because they are widely used. We include a comparison to BWA-MEM [14] for the same reason. While we did not modify the newer HISAT2 (http://ccb.jhu.edu/software/hisat2), the same modifications should benefit that software as well. We ran HISAT with the `--no-spliced-alignment --no-temp-splicesite` options to disable gathering of splice-site evidence because our input reads were from DNA sequencing experiments. Software used to run the experiments and produce the figures and tables is located at https://github.com/BenLangmead/bowtie-scaling.

### Reads

We obtained sequencing reads from accessions ERR194147 (Platinum Genomes Project [29]), SRR069520 (1000 Genomes Project [30]) and SRR3947551 (a low coverage whole genome sequencing project [31]). All reads are 100 × 100 nt (paired-end) from the Illumina HiSeq 2000 instrument. We downloaded the reads in FASTQ format, selected a random subset of 100M from each of the 3 accessions, then randomized the order of the resulting set of 300M reads to avoid clustering of reads with similar properties. These constitute the *human_100_300M* input read set. Bowtie is designed to align shorter reads, so we also created set *human_50_300M* consisting of the *human_100_300M* reads truncated to 50 nt at the 3’ end. Unpaired alignment experiments use just the first-end FASTQ files. Download links these reads are in Supplementary Note 2.

### Evaluation systems

Each job was run on three servers, which we call *Broadwell*, *Skylake* and *KNL* for short. Broadwell is a dual-socket system with two Intel Xeon E7-4830 v4 2.00GHz CPUs and 1 TB of DDR4 memory. Both CPUs have 28 physical processor cores, enabling up to 112 threads of execution since each core supports 2 simultaneous “hyperthreaded” threads. The system runs CentOS 6.8 Linux, kernel v2.6.32, and is located at the Maryland Advanced Research Computing Center (MARCC). Skylake is an Intel Xeon Platinum 8160 system with 192GB of memory and 48 physical processor cores, enabling up to 96 threads of execution, 2 per core. This system runs CentOS Linux release 7.4.1708, kernel v3.10.0, and is located in the Stampede 2 cluster at the Texas Advanced Compute Center (TACC) accessible via the XSEDE network. KNL is an Intel Xeon Phi 7250 (Knight’s Landing) system with 96GB DDR4 memory (as well as a 16GB high-speed MCDRAM). The system has 68 physical processor cores, enabling up to 272 threads of execution, 4 per core. This system is also located in the Stampede 2 cluster and the operating system and kernel are identical to the Skylake system.

Although these three platforms differ in architectural details - e.g. in the number of simultaneous threads allowed - we test the same parallelization schemes on all three. Since the three systems support the same basic instruction set, we are running exactly the same executables on all three.

In all experiments, FASTQ input is read from a local disk and SAM output is written to the same local disk. In the case of the Broadwell system, the disk is a magnetic 7200 RPM SATA hard drive. In the case of Skylake and KNL, the disk is a local solid-state drive.

### 3.1 Varying lock type

As discussed in Methods, we extended Bowtie, Bowtie 2 and HISAT to use one of four lock types: a `TinyThread++` spinlock, TBB standard lock, TBB spinlock, or TBB queueing lock. For each run, we launched a single aligner process configured for multithreading (MT), using the `-p` option to specify the number of simultaneous threads, *T*. The MP baseline used the TBB queueing lock. When plotting, we arranged thread count on the horizontal axis and maximum per-thread wall-clock time (i.e. time required to align all reads) on the vertical axis. Because we vary the number of input reads in direct proportion to thread count, ideal scaling would show as a flat horizontal line, whereas worse-than-ideal scaling shows as an upward-trending line. Since we omit runs that took over 20 minutes, some lines “fall off” the top of the plot.

Figure 3 shows how thread count affects running time for unpaired alignment. Supplementary Figure 1 shows the same for paired-end alignment. We observe that the MP baseline outperformed all multithreading modes (MT). Choice of lock type clearly impacts scaling, seen mostly clearly in the Bowtie and HISAT configurations. While no lock type performed best in all cases, the TBB queueing lock tended to eventually outperform other MT configurations at high thread count. This is clearest for HISAT and Bowtie. There were also cases where the TBB standard lock outperformed the queueing lock at the very highest thread counts, as seen in the Skylake+HISAT, KNL+Bowtie 2 and Broadwell+Bowtie results.

**Figure 3:**
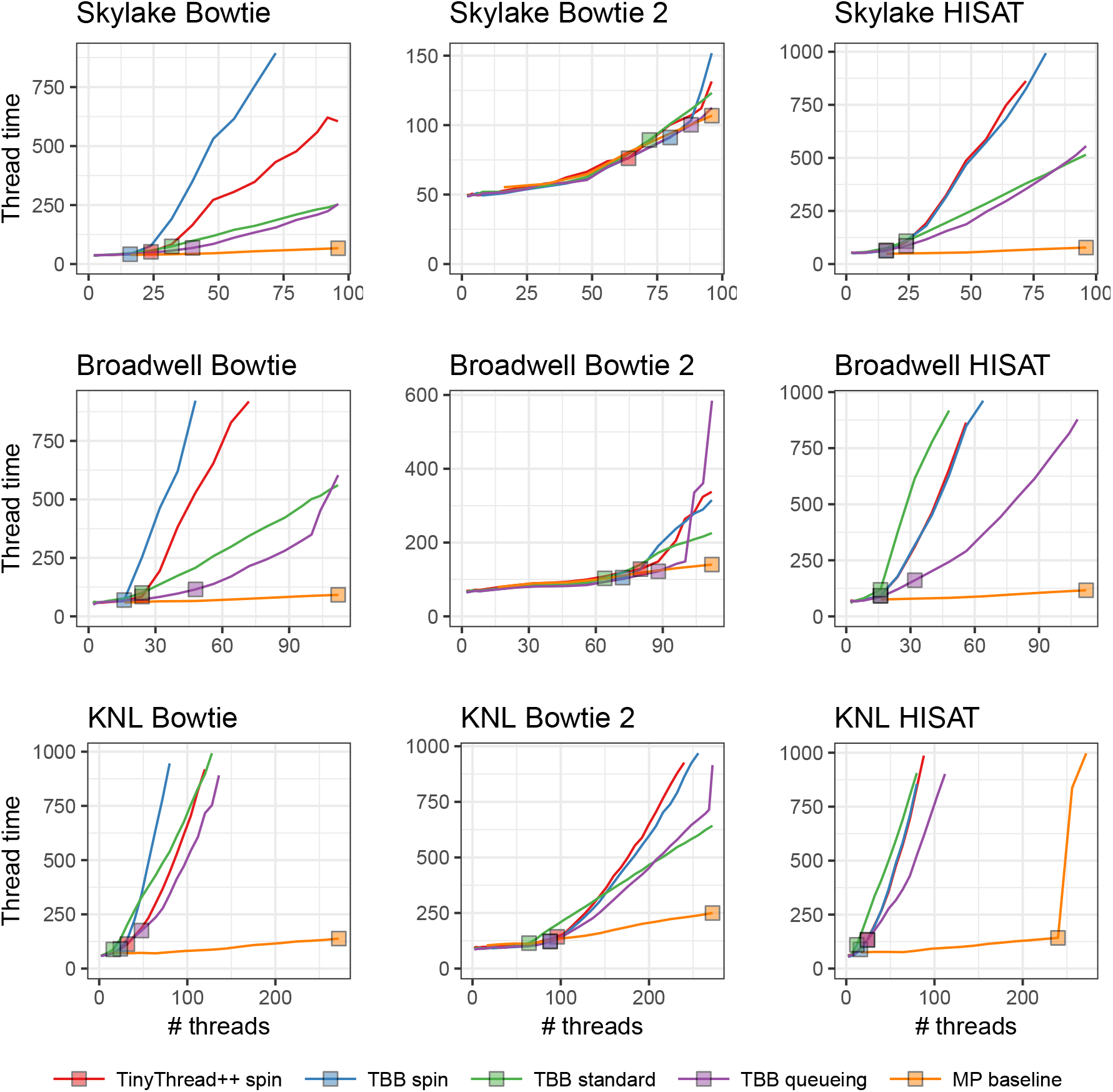
Comparison of 4 lock types and multiprocessing baseline. Reads are unpaired. Results are shown for three aligners (rows) and three systems (columns). Jobs that ran for over 20 minutes are omitted. Squares indicate the point on each line yielding maximal total alignment throughput. These points are summarized in Table 2.

Table 2 shows peak throughputs (also represented by squares in Figure 3) for each lock type and the MP baseline. For 11 out of 18 combinations of aligner, test system and paired-end status, the queueing lock has the second-highest peak throughput after the MP baseline.

**Table 2.** Peak throughputs for four lock types and multiprocessing baseline. For each row, maximal peak throughput in thousands of reads per second (Krd/s) and number of threads that achieved the peak (Th) are reported. For each combination of aligner, paired-end status and test system, the best and second-best throughputs are highlighted red and orange respectively.

|  |  | Skylake (96 threads) |  |  |  | Broadwell (112 threads) |  |  |  | KNL (272 threads) |  |  |  |
| --- | --- | --- | --- | --- | --- | --- | --- | --- | --- | --- | --- | --- | --- |
|  |  | Paired |  | Unpaired |  | Paired |  | Unpaired |  | Paired |  | Unpaired |  |
|  |  | Th | Krd/s | Th | Krd/s | Th | Krd/s | Th | Krd/s | Th | Krd/s | Th | Krd/s |
| Bowtie | TinyThread++ spin | 92 | 126.54 | 24 | 446.94 | 88 | 98.69 | 24 | 282.93 | 72 | 41.96 | 32 | 127.48 |
|  | TBB spin | 96 | 130.98 | 16 | 378.96 | 88 | 97.01 | 16 | 229.62 | 72 | 42.92 | 24 | 118.74 |
|  | TBB standard | 80 | 121.79 | 32 | 405.59 | 72 | 86.08 | 24 | 239.80 | 48 | 26.54 | 16 | 81.67 |
|  | TBB queueing | 92 | 132.38 | 40 | 586.60 | 96 | 103.61 | 48 | 415.78 | 72 | 39.22 | 48 | 123.33 |
|  | MP baseline | 96 | 128.66 | 96 | 1,433.82 | 112 | 108.89 | 112 | 1,215.75 | 272 | 70.44 | 272 | 895.90 |
| Bowtie 2 | TinyThread++ spin | 80 | 65.55 | 72 | 176.27 | 80 | 51.55 | 80 | 125.02 | 96 | 16.49 | 96 | 43.71 |
|  | TBB spin | 88 | 68.02 | 80 | 177.78 | 64 | 55.45 | 72 | 137.07 | 96 | 18.17 | 88 | 47.09 |
|  | TBB standard | 56 | 62.47 | 64 | 162.36 | 56 | 49.05 | 64 | 124.19 | 64 | 12.73 | 64 | 36.36 |
|  | TBB queueing | 80 | 66.60 | 88 | 180.83 | 64 | 54.48 | 88 | 144.14 | 96 | 17.72 | 88 | 46.64 |
|  | MP baseline | 96 | 67.62 | 96 | 185.25 | 112 | 57.11 | 112 | 159.47 | 272 | 27.98 | 272 | 69.52 |
| HISAT | TinyThread++ spin | 16 | 134.13 | 16 | 292.61 | 16 | 89.80 | 16 | 210.14 | 16 | 29.11 | 24 | 71.72 |
|  | TBB spin | 16 | 136.87 | 16 | 307.35 | 16 | 94.10 | 16 | 209.14 | 16 | 29.55 | 16 | 72.18 |
|  | TBB standard | 16 | 99.28 | 16 | 250.40 | 16 | 61.89 | 16 | 162.16 | 8 | 19.96 | 12 | 43.60 |
|  | TBB queueing | 16 | 137.84 | 24 | 349.36 | 16 | 98.45 | 32 | 238.34 | 32 | 27.65 | 24 | 72.56 |
|  | MP baseline | 96 | 710.58 | 96 | 1,478.72 | 112 | 568.00 | 112 | 1,154.60 | 272 | 360.41 | 240 | 686.50 |

In some cases, queueing lock performance deteriorated quickly at the highest thread counts, e.g. for Broadwell+Bowtie, Broadwell+Bowtie 2 and KNL+Bowtie 2. This contrasts with the TBB standard lock, which deteriorated more slowly (and almost linearly) at high thread counts. This is likely due to starvation; at high thread counts, threads share cores (up to 2 threads per cores on Broadwell, 4 on KNL), so the optimistic queueing lock will tend to spin fruitlessly on contended locks, starving the lock holder. This problem is not shared by the pessimistic standard lock.

Even the best-scaling configuration — the MP baseline — had less than perfect scaling. Increasing thread count increases contention for shared resources, slowing all threads on average. For example, higher thread count leads to greater contention for shared memory, e.g. L1 and L2 caches, translation look-aside buffer, and arithmetic and vector processing units. This is more obvious on the KNL system where up to 4 threads can share a processor.

Divergence between the lock-type scaling behaviors was lower for Bowtie 2 than for the other tools. This is likely because Bowtie 2 requires more time to align a single read. This spreads locking attempts out over time and thereby reduces contention. Thus, Bowtie 2’s lower divergence is consistent with the theory that differences come primarily from contention overhead.

In summary: while the MP baseline outperformed all MT configurations, the TBB queueing lock often scaled best, with the TBB standard lock doing well or better in some situations. Supplementary Table 2 shows how these peak throughputs translate to wall-clock time required to align 100 nt reads covering the human genome to 40-fold average depth; e.g. in the case of paired-end KNL+Bowtie 2, moving from the TBB spin lock to the queueing lock reduces running time from about 26 hours to about 19 hours. We use the queueing lock in subsequent Bowtie, Bowtie 2 and HISAT experiments.

### 3.2 Varying parsing method

The gap between MP baseline and MT methods spurred us to examine input and output synchronization. We hypothesized the gap was due to a combination of (a) length of time spent in these critical sections, and (b) overhead of locking and unlocking. We tried to close the gap using the strategies discussed in Methods: deferred (D-) parsing and batch (B-) parsing. B-parsing used a batch size of 32 in all experiments. Figure 4 shows running time versus thread count for unpaired alignment using each strategy. Supplementary Figure 2 shows the same for paired-end alignment. A clear ordering exists among the strategies: B-parsing outperformed D-parsing, which outperformed O-parsing. This was basically true in every scenario tested. B-parsing scaled well enough to be competitive with the MP baseline in multiple scenarios, e.g. for Bowtie 2 and for all paired-end Bowtie and Bowtie 2 scenarios.

**Figure 4:**
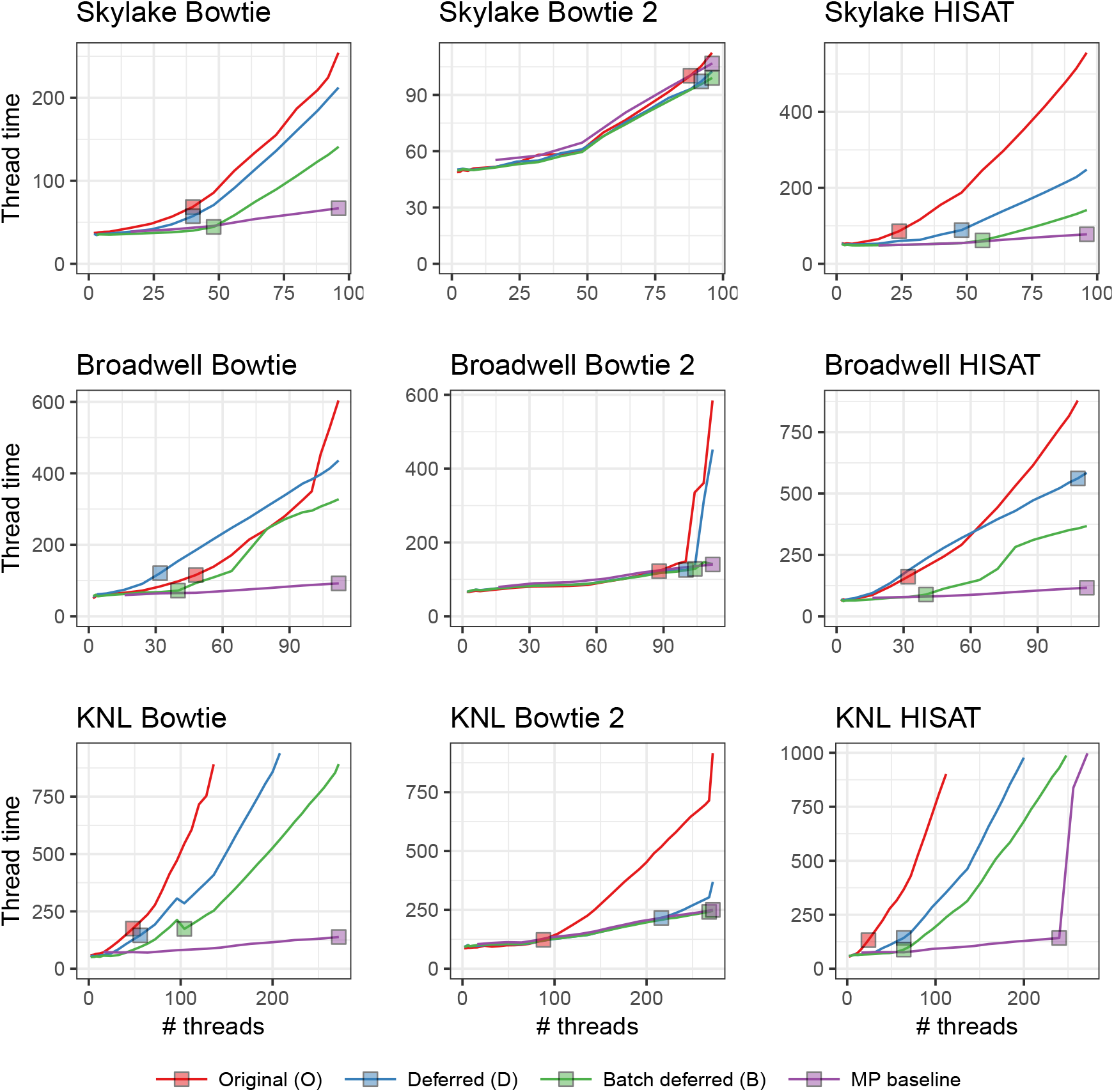
Comparison of 3 parsing strategies and multiprocessing baseline. Reads are unpaired. Jobs that ran for over 20 minutes are omitted. Squares indicate the point on each line yielding maximal total alignment throughput and these points are summarized in Table 3.

Table 3 shows peak throughputs (represented by squares in Figure 4) for each strategy the MP baseline. B-parsing had either the highest or second-highest peak throughput in all scenarios except paired-end KNL+Bowtie 2, where it slightly underperformed D-parsing. Moving from O-parsing to B-parsing for unpaired KNL+Bowtie 2 reduces extrapolated human-40x-coverage running time from about 7h:10m to about 4h:40m, bringing it below BWA-MEM’s 6h:40m running time (Supplementary Table 2).

**Table 3:** Peak throughputs for three parsing strategies and multiprocessing baseline. For each combination of aligner, paired-end status and test system, the best and second-best throughputs are highlighted red and orange respectively. The TBB queueing lock is used in all cases.

|  |  | Skylake (96 threads) |  |  |  | Broadwell (112 threads) |  |  |  | KNL (272 threads) |  |  |  |
| --- | --- | --- | --- | --- | --- | --- | --- | --- | --- | --- | --- | --- | --- |
|  |  | Paired |  | Unpaired |  | Paired |  | Unpaired |  | Paired |  | Unpaired |  |
|  |  | Th | Krd/s | Th | Krd/s | Th | Krd/s | Th | Krd/s | Th | Krd/s | Th | Krd/s |
| Bowtie | O-parsing | 92 | 132.38 | 40 | 586.60 | 96 | 103.61 | 48 | 415.78 | 72 | 39.22 | 48 | 123.33 |
|  | D-parsing | 96 | 141.48 | 32 | 563.14 | 104 | 102.51 | 32 | 264.94 | 136 | 58.59 | 32 | 170.60 |
|  | B-parsing | 96 | 146.47 | 56 | 1,257.81 | 112 | 115.41 | 40 | 556.38 | 200 | 70.99 | 40 | 263.08 |
|  | MP baseline | 96 | 128.66 | 96 | 1,433.82 | 112 | 108.89 | 112 | 1,215.75 | 272 | 70.44 | 272 | 895.90 |
| Bowtie 2 | O-parsing | 80 | 66.60 | 88 | 180.83 | 64 | 54.48 | 88 | 144.14 | 96 | 17.72 | 88 | 46.64 |
|  | D-parsing | 92 | 71.63 | 92 | 191.14 | 104 | 58.93 | 100 | 159.18 | 268 | 27.51 | 224 | 67.68 |
|  | B-parsing | 96 | 72.00 | 96 | 199.55 | 104 | 58.41 | 104 | 161.77 | 272 | 26.61 | 268 | 71.13 |
|  | MP baseline | 96 | 67.62 | 96 | 185.25 | 112 | 57.11 | 112 | 159.47 | 272 | 27.98 | 272 | 69.52 |
| HISAT | O-parsing | 16 | 137.84 | 24 | 349.36 | 16 | 98.45 | 32 | 238.34 | 32 | 27.65 | 24 | 72.56 |
|  | D-parsing | 40 | 285.80 | 32 | 497.85 | 108 | 144.26 | 108 | 230.70 | 32 | 69.12 | 40 | 146.71 |
|  | B-parsing | 56 | 522.93 | 48 | 1,016.27 | 56 | 315.24 | 40 | 540.92 | 48 | 116.58 | 40 | 215.73 |
|  | MP baseline | 96 | 710.58 | 96 | 1,478.72 | 112 | 568.00 | 112 | 1,154.60 | 272 | 360.41 | 240 | 686.50 |

The MP baseline had the highest peak throughput in 10 of 18 scenarios, including all the HISAT scenarios. There was still a wide gap between the MP baseline and the best-performing MT configuration in many scenarios, particularly for Bowtie and HISAT. As when investigating lock type, we found divergence between parsing strategies was lower for Bowtie 2 than for the other tools. This is likely because Bowtie 2 spent more time aligning each read compared to the others, reducing contention.

### 3.3 Final evaluations

Finally we compared B-parsing to block deferred (L-) parsing. L-parsing’s critical section is the simplest, so we hypothesized it would outperform B-parsing. But since L-parsing requires padded input, using it in practice requires an initial pass to add the padding, which might itself become the bottleneck. We revisit L-parsing’s practicality in the Discussion section.

To test block parsing (L) we created padded input sets (Figure 2). We created one new set called *human_100_block_300M* with the same reads as *human_100_300M* but padding the FASTQ to achieve 12 KB blocks (*B* = 12288) and 44 reads per block (*N* = 44). Similarly, we created a set called *human_50_block_300M* with the reads from *human_50_300M* padded to achieve 12 KB blocks (*B* = 12288) and 70 reads per block (*N* = 70). *N* and *B* are specified to the aligner via command-line options (`--block-bytes` and `--reads-per-block`).

We tested two versions of L-parsing, one that writes output to a single SAM file and one that stripes output across 16 SAM files, with each thread writing to an output file corresponding to the thread ID modulo 16. The striped output mode was added after noticing poor performance due to output lock contention at high thread counts on KNL (Supplementary Figure 3).

For Bowtie 2, we also compared to BWA-MEM v0.7.16a [14] with default arguments. BWA-MEM uses a pipelined multithreading strategy. Two master threads run simultaneously, each cycling through three steps: (a) parsing a batch of input reads, (b) aligning the batch, and (c) writing the output alignments for the batch. Using a pthreads [32] lock and condition variable, the master threads are prevented from running the aligning step at the same time; when one thread is aligning, the other is writing output or reading input. When in the alignment step, the master thread spawns *T* worker threads, *T* given by the `-t` option. Worker threads balance load using work stealing, synchronizing with atomic operations. Batch size is determined by multiplying a number of input bases (10 million) by *T*. We note that (a) these are large batches compared to Bowtie 2, which uses a batch size of 32 reads for B-parsing and at most 70 for L-parsing, and (b) that, while the batch size is independent of thread count for Bowtie 2, it grows linearly with thread count in BWA-MEM. BWA-MEM experiments used the same number of input reads as the Bowtie 2 experiments (Supplementary Table 1). We also corrected an issue in the BWA-MEM code that caused failures for thread counts over 214, a limit we exceed on KNL (Supplementary Note 3).

Figure 5 shows the comparison for unpaired alignment and Supplementary Figure 4 shows the same for paired-end alignment. Table 4 gives maximal peak throughput for each configuration. While L- and B-parsing scaled similarly at low thread counts, L-parsing maintained excellent scaling through higher thread counts in most configurations. L-parsing with striped output was either best or competitive for all scenarios. B-parsing scaled substantially worse than L-parsing for HISAT and for unpaired Bowtie.

**Figure 5:**
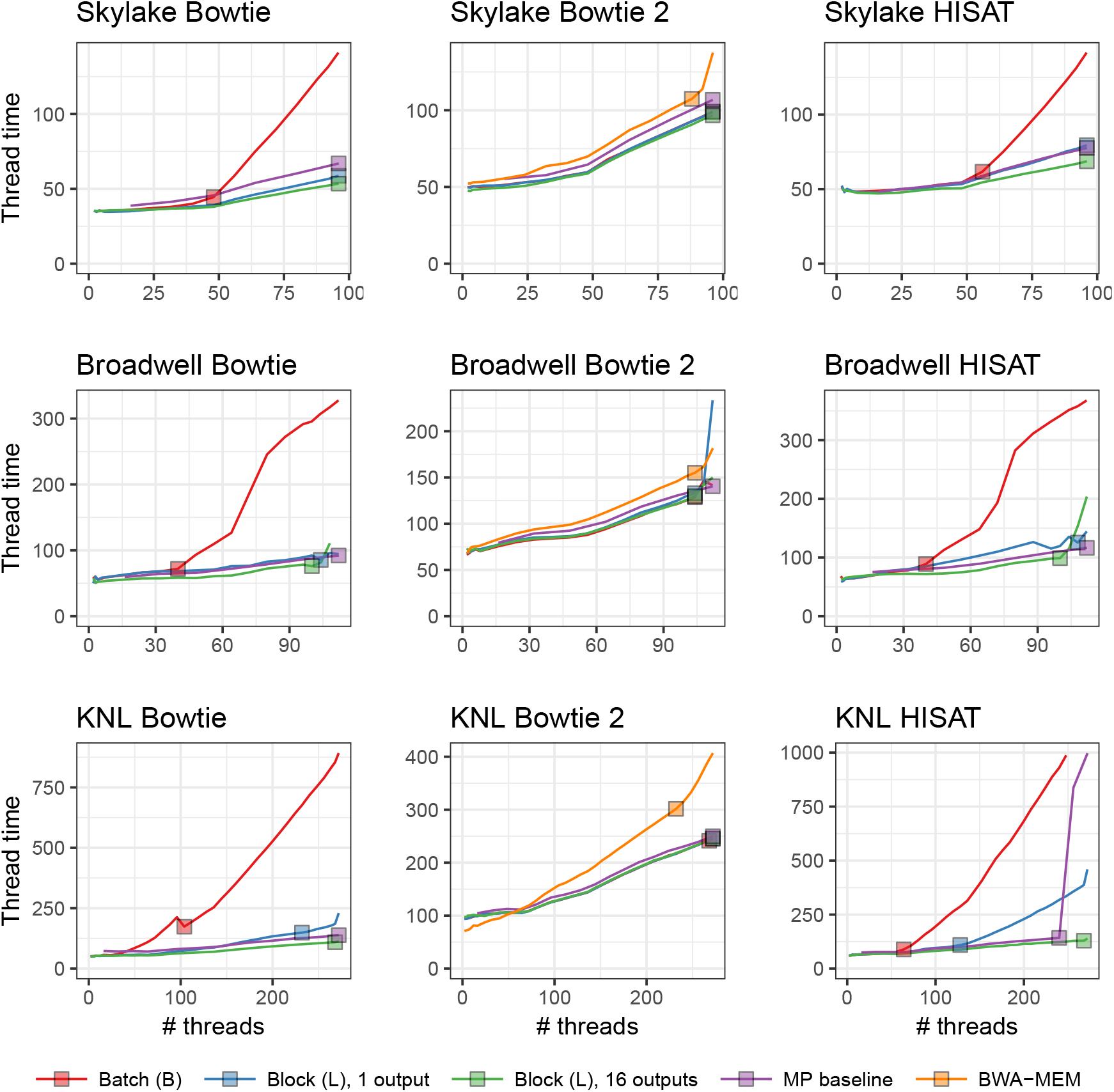
Unpaired-alignment comparison of B-parsing, L-parsing, L-paring with output striped across 16 files and the MP baseline. BWA-MEM is also evaluated and compared to the Bowtie 2 configurations. Jobs that ran for over 20 minutes are omitted. Squares indicate the run for each configuration yielding greatest overall alignment throughput, also summarized in Table 4.

**Table 4.**
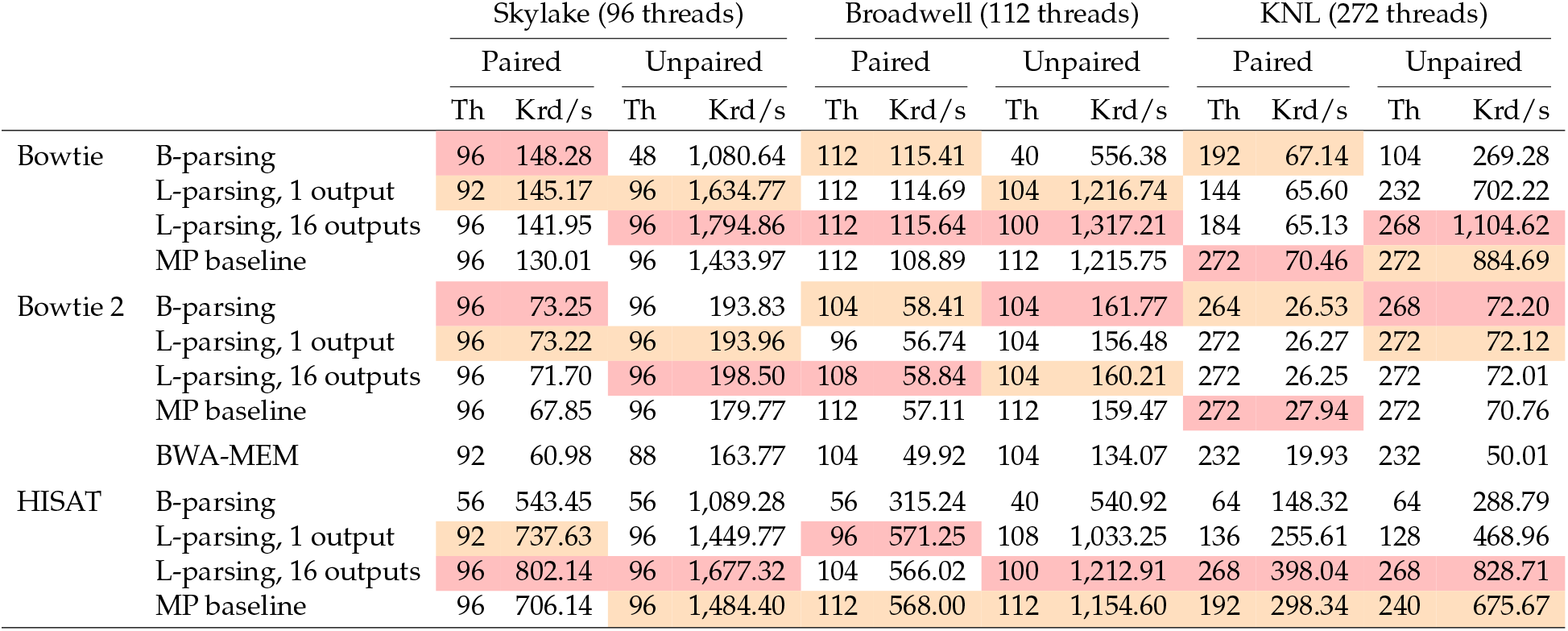
Peak throughputs for B-parsing, L-parsing, L-paring with output striped across 16 files, and the MP baseline. BWA-MEM is also evaluated for the Bowtie 2 configurations. For each combination of aligner, paired-end status and test machine, the best and second-best throughputs are highlighted in red and orange respectively.

|  |  | Skylake (96 threads) |  |  |  | Broadwell (112 threads) |  |  |  | KNL (272 threads) |  |  |  |
| --- | --- | --- | --- | --- | --- | --- | --- | --- | --- | --- | --- | --- | --- |
|  |  | Paired |  | Unpaired |  | Paired |  | Unpaired |  | Paired |  | Unpaired |  |
|  |  | Th | Krd/s | Th | Krd/s | Th | Krd/s | Th | Krd/s | Th | Krd/s | Th | Krd/s |
| Bowtie | B-parsing | 96 | 148.28 | 48 | 1,080.64 | 112 | 115.41 | 40 | 556.38 | 192 | 67.14 | 104 | 269.28 |
|  | L-parsing, 1 output | 92 | 145.17 | 96 | 1,634.77 | 112 | 114.69 | 104 | 1,216.74 | 144 | 65.60 | 232 | 702.22 |
|  | L-parsing, 16 outputs | 96 | 141.95 | 96 | 1,794.86 | 112 | 115.64 | 100 | 1,317.21 | 184 | 65.13 | 268 | 1,104.62 |
|  | MP baseline | 96 | 130.01 | 96 | 1,433.97 | 112 | 108.89 | 112 | 1,215.75 | 272 | 70.46 | 272 | 884.69 |
| Bowtie 2 | B-parsing | 96 | 73.25 | 96 | 193.83 | 104 | 58.41 | 104 | 161.77 | 264 | 26.53 | 268 | 72.20 |
|  | L-parsing, 1 output | 96 | 73.22 | 96 | 193.96 | 96 | 56.74 | 104 | 156.48 | 272 | 26.27 | 272 | 72.12 |
|  | L-parsing, 16 outputs | 96 | 71.70 | 96 | 198.50 | 108 | 58.84 | 104 | 160.21 | 272 | 26.25 | 272 | 72.01 |
|  | MP baseline | 96 | 67.85 | 96 | 179.77 | 112 | 57.11 | 112 | 159.47 | 272 | 27.94 | 272 | 70.76 |
|  | BWA-MEM | 92 | 60.98 | 88 | 163.77 | 104 | 49.92 | 104 | 134.07 | 232 | 19.93 | 232 | 50.01 |
| HISAT | B-parsing | 56 | 543.45 | 56 | 1,089.28 | 56 | 315.24 | 40 | 540.92 | 64 | 148.32 | 64 | 288.79 |
|  | L-parsing, 1 output | 92 | 737.63 | 96 | 1,449.77 | 96 | 571.25 | 108 | 1,033.25 | 136 | 255.61 | 128 | 468.96 |
|  | L-parsing, 16 outputs | 96 | 802.14 | 96 | 1,677.32 | 104 | 566.02 | 100 | 1,212.91 | 268 | 398.04 | 268 | 828.71 |
|  | MP baseline | 96 | 706.14 | 96 | 1,484.40 | 112 | 568.00 | 112 | 1,154.60 | 192 | 298.34 | 240 | 675.67 |

Remarkably, L-parsing with striped output scaled better than the MP baseline in all but a few cases, and best overall in 11 out of 18 cases. Thus, L-parsing with striped output is the only approach we evaluated that improved on the MP baseline.

The 1-output and 16-output (striped) versions of L-parsing scale similarly with the notable exception of KNL+HISAT, where the 1-output versions scales substantially worse. This comports with the fact that HISAT is the fastest of the aligners tested and therefore generates output the most rapidly. This causes increased contention for the output lock in the 1-output version. In the 16-output version, the load is spread over 16 locks, reducing overall contention.

While BWA-MEM scaled well, both the B-parsing and L-parsing Bowtie 2 configurations scaled better. This was particularly true on the KNL system, where B-parsing achieved 33% (paired-end) and 45% (unpaired) higher throughput and L-parsing achieved 32% (paired-end) and 44% (unpaired) higher throughput. When translated to extrapolated 40x-human running time, Bowtie 2’s unpaired B-parsing mode finishes about 2 hours faster than BWA-MEM, and its paired-end B-parsing mode finishes about 4 hours faster (Supplementary Table 2). BWA-MEM’s larger input chunk size, together with the chunk size’s linear scaling, also caused BWA-MEM’s memory footprint to grow much faster than Bowtie 2’s (Supplementary Figure 5).

## 4 Discussion

General-purpose processors now support hundreds of simultaneous threads of execution and future architectures will likely continue the trend of squeezing more relatively slow threads onto a single chip. Genomics software must adapt to high thread counts, slow individual threads, and system architectures that more closely resemble small computer clusters — complete with interconnection network and distributed storage — than simpler processors of the past.

We addressed how lock types, design of critical sections, NUMA, starvation and other issues can impact thread scaling on two Intel systems, including one based on the many-core Knight’s Landing architecture. We greatly improved thread scaling for three commonly used alignment tools: Bowtie, Bowtie 2 and HISAT. We measured the effect of each candidate improvement, and also showed that the improvements to Bowtie 2 allow it to scale more favorably than BWA-MEM with respect to both time and peak memory footprint. The TBB queueing lock and the B-parsing method are the default as of Bowtie v1.2.0 and Bowtie 2 v2.3.0.

Bowtie and HISAT align reads more quickly than Bowtie 2 (Table 4), making their locks more contended and thread scaling more difficult. This is reinforced by how much L-parsing improved thread scaling for Bowtie and HISAT. This suggests that similar or greater gains may be possible by adapting our methods to yet faster tools such as pseudoaligners [33], quasi-mappers [34] and tools that analyze at the *k*-mer level [13].

There is also further room for improvement. Besides L-parsing, which requires special padding, B-parsing was the best-scaling MT strategy and it is now the default strategy in Bowtie and Bowtie 2. But the MP baseline outperformed B-parsing in some scenarios, and L-parsing outperformed it nearly always. We still seek MT methods that scale like L-parsing but that work with standard, unpadded inputs.

For further gains, it will be important to investigate more lock types. The queueing (MCS) lock [26] scaled best at high thread counts, likely because of reduced cache-coherence communication. But other lock types could improve on this in two ways. First: like the spinlock, the queueing lock is optimistic. But when thread count and contention are high, pessimism is more appropriate. It will be important to investigate lock types that adapt their degree of optimism in inverse proportion to the lock contention, e.g. the hierarchical backoff lock [35]. Secondly, while the queueing lock successfully reduces cache-coherence communication, other locks go further in this regard. The cohort lock [36] further reduces communication by maximizing the chance that consecutive holders of the lock are physically proximate (i.e. on the same NUMA node), avoiding longer-distance communication.

Genomics file formats, notably FASTQ and FASTA, have properties that impede thread scaling. Since records lack predictable length, record boundaries must be identified in a synchronized manner, i.e. inside a critical section. Our best scaling results were achieved by forcing predictable FASTQ record boundaries (using padding) and simplifying the input critical section to a single fixed-size read. This padding is easy to add, regardless of the reads’ paired-end status or length (including mixed lengths within a file and between paired ends) as long as the block size *B* accommodates the longest read. While this suggests a strategy of pre-padding FASTQ files prior to L-parsing alignment, that might simply move the synchronization bottleneck into the padding step. It may be worth the cost, though, if input files are to be re-used across multiple L-parsing alignment jobs, amortizing the padding cost.

L-parsing padding consists of simple runs of space characters, which are highly compressible. For our inputs, padding increased uncompressed FASTQ file size by 9-14%, but gzipped FASTQ file size increased just 1.0-1.5% (Supplementary Table 3). More generally, it is common to store sequencing reads in a compressed form, then decompress — e.g. with gzip or the libz library — prior to read alignment. But if decompression must be performed either upstream of the read aligner or in the aligner’s input critical section, decompression is liable to become a new thread-scaling bottleneck. A possible workaround is similar to the idea behind D-parsing: instead of both reading and decompressing in the critical section, decompression could be deferred until after the critical section. This might be facilitated by block-compressed formats like BGZIP [37]. A question for future work is whether compressed inputs can be used to reduce the space overhead of padding while still providing thread scaling similar to what we achieved here with L-parsing and uncompressed inputs. A related question is whether compressed output would mitigate bottlenecks of the kind we (mostly) avoided with our multiple-output-file scheme (Supplementary Figure 3).

Finally, we note that the threading model we consider here, whereby all threads rotate through input, alignment and output phases (Figure 1), is just one possible model. We described the alternative model used in BWA-MEM in Section 3.3. Another popular model is to relegate input parsing and output writing to separate specialized threads. The input thread only parses input, placing parsed records onto a queue to be later retrieved by alignment threads. Because only one thread reads input, no locking is needed. Similarly, the output thread only writes output (without locking), receiving alignment records from a queue populated by alignment threads. Synchronization is still required, but it is limited to points where items are added to or removed from queues. The tools examined here can be adapted to use this alternate model, and we already did so for Bowtie 1’s unpaired alignment mode. Results are mixed but promising: a branch of Bowtie’s B-parsing code that relegates input and output to separate threads achieves better thread scaling on KNL, but worse on Skylake compared to standard B-parsing (Supplementary Figure 6). It will be important to compare and contrast such threading models, and to test how they interact with compressed inputs and outputs, in future work.

## 5 Acknowledgments

We are grateful to many at Intel and the Parallel Computing Center program, for technical and administrative assistance, including John Oneill, Ram Ramanujam, Kevin O’leary, Lisa Smith and Brian Napier. This work used the Extreme Science and Engineering Discovery Environment (XSEDE), supported by National Science Foundation grant number ACI-1548562.

